# Transcriptomic Profiling of Plasma Extracellular Vesicles Enables Reliable Annotation of the Cancer-specific Transcriptome and Molecular Subtype

**DOI:** 10.1101/2022.10.27.514047

**Authors:** Vahid Bahrambeigi, Jaewon J. Lee, Vittorio Branchi, Kimal I. Rajapakshe, Zhichao Xu, Jason T. Henry, Wang Kun, Bret M. Stephens, Sarah Dhebat, Mark W. Hurd, Ryan Sun, Peng Yang, Eytan Ruppin, Wenyi Wang, Scott Kopetz, Anirban Maitra, Paola A. Guerrero

## Abstract

Longitudinal monitoring of patients with advanced cancers is crucial to evaluate both disease burden and treatment response. Current liquid biopsy approaches mostly rely on the detection of DNA-based biomarkers. However, plasma RNA analysis can unleash tremendous opportunities for tumor state interrogation and molecular subtyping. Through the application of deep learning algorithms to the deconvolved transcriptomes of RNA within plasma extracellular vesicles (evRNA), we successfully predict consensus molecular subtypes in metastatic colorectal cancer patients. We further demonstrate the ability to monitor changes in transcriptomic subtype under treatment selection pressure and identify molecular pathways in evRNA associated with recurrence. Our approach also identified expressed gene fusions and neoepitopes from evRNA. These results demonstrate the feasibility of transcriptomic-based liquid biopsy platforms for precision oncology approaches, spanning from the longitudinal monitoring of tumor subtype changes to identification of expressed fusions and neoantigens as cancer-specific therapeutic targets, *sans* the need for tissue-based sampling.

**Statement of significance:** We have developed an approach to interrogate changes in cancer molecular subtypes and differentially expressed genes, through the analysis and deconvolution of RNA sequencing of plasma EVs. Serial analyses of tumor-encoded transcriptomes in liquid biopsies can enable facile cancer detection and monitor for recurrences and therapy-induced tumor evolution.

## Introduction

Liquid biopsies can detect cancer-derived materials without the need to perform an invasive tissue biopsy. Most liquid biopsy assays designed for therapeutic assignment and response monitoring in cancer are currently based on circulating cell-free DNA (cfDNA), such as the detection of mutations, fragmentation patterns or methylation signatures [1-4]. Recent studies have made initial forays into inferring tumor transcriptomes indirectly via the use of fragmented cfDNA profiles in plasma samples, and identifying pertinent biological pathways based on the anatomical location of the primary cancer [5, 6]. Nonetheless, the direct interrogation of tumor transcriptomes in plasma could enable significant insights into the cancer molecular landscape beyond what genomic alterations alone would provide. The comprehensive assessment of cancer-derived RNA in plasma has been challenging due to the presence of circulating ribonucleases, which cleave coding RNAs, confounding their assessment with next generation sequencing (NGS) platforms, and restricting most studies of this nature to the non-coding transcriptome [7, 8]. A second major challenge of a circulating RNA-based approach is the reliable separation of tumor-derived transcriptomic signatures from the “background” RNA released by non-neoplastic cell types. Among liquid biopsy substrates, extracellular vesicles (EVs) represent a facile compartment for querying tumor transcriptomes as they are actively released from cancer cells and contain intact RNA cargoes (evRNA) [9]. While we and others have previously demonstrated the ability to map the DNA landscapes of advanced cancers reliably with EVs [10, 11], the feasible extrapolation of this to cancer-derived evRNA has not been documented.

In this paper, we describe our pipeline for isolating, sequencing, and interrogating the whole transcriptomes of advanced colorectal cancer (CRC) using plasma EVs. Notably, the use of evRNA cargo overcomes the current limitation of sequencing extensively fragmented RNAs in circulation, while the application of machine learning facilitates deconvolution of cancer-specific transcriptome data from the cumulative “pool” of plasma EVs. We demonstrate our ability to assign current widely accepted RNA-based tumor subtype signatures typically obtained via tissue samples – namely, Consensus Molecular Subtypes (CMS) for CRC in baseline plasma samples [12]. We further demonstrate the ability of our pipeline to longitudinally monitor changes in evRNA-based CMS subtype and differential gene expression under treatment selection, which was clinically associated with disease progression. While such “subtype switching” as a basis for treatment resistance is well established [13], it is based entirely on serial sampling of tumor tissue, a requirement that can now be precluded with longitudinal monitoring of evRNAs. Finally, our evRNA-based profiling identifies expressed gene fusions and neoantigens in the circulation of CRC patients, which has significant implications for monitoring those on targeted and immunotherapies. The pipeline described herein should be applicable across cancer types and provide an avenue for leveraging the full potential of mapping tumor transcriptomes and their evolution over time, *sans* the requirement for repeated tissue biopsies.

## Results

### EVs transcriptomic profiles recapitulate the transcriptomic profiles of the cells of origin in vitro

It is well established that cancer cell-derived EVs contain mRNA cargo [14-16]. However, the correlation between cellular RNA and the derivative evRNA with respect to transcriptome-based cancer subtyping has not been explored. To address this question, we performed RNA sequencing (RNA-seq) on total cellular RNA (cellRNA) and evRNA isolated from different CRC cell lines, which were previously classified according to the four-tier CMS classification system (CMS1-4, respectively) [17, 18]. We first applied CMScaller, a CMS classifier optimized to stratify samples for both *in vitro* and *in vivo* models [19, 20]. Utilizing a customized panel of CMS template genes, the expression profiles of CMS genes were comparable between matched cellRNA and evRNA for individual cell lines (**Figure S1a**). In addition, utilizing a previously described machine learning approach to classify molecular subtypes [21], we observed a 100% CMS concordance between EVs and corresponding cells of origin across all cell lines (**Figures S1b**).

### A deconvolution pipeline identifies tumor-specific evRNA in plasma

Plasma samples contain both tumor and non-tumor derived EVs mixed in unknown proportions. To overcome this intrinsic limitation of the evRNA analyte, we designed a series of experiments to bioinformatically deconvolve evRNA into tumor and non-tumor-derived transcripts, and then classify the tumor-derived transcripts [21] (**Figure S1c-d**). As a proof of concept, we applied a computational deconvolution pipeline on *in vitro* cellRNA mixtures containing known proportions of cancer and normal RNAs to infer cancer-derived RNA proportions and test whether the cancer-specific transcriptome can be used for downstream analyses. For *in vitro* cellRNA mixtures, we mixed extracted RNA from a CRC cell line (one cell line per CMS) with RNA from a normal colon cell line (841-CoN) at different rations ranging from 0.1% to 50% cancer-derived RNA (**Figure S1c**). Using CIBERSORTx, we generated a signature matrix consisting of genes that can discriminate each compartment of interest (cancer and non-cancer) to impute cancer proportion and expression signatures [22, 23] (**Figure 1a, Figure S1c-d**). In addition to CIBERSORTx, we applied a recently developed tool, named the COnfident DEconvolution For All Cell Subsets (CODEFACS) (**Figure 1a**) [24]. The signature matrix and imputed proportions generated by CIBERSORTx along with cellRNA mixtures were used as input for CODEFACS. After data deconvolution, we classified the deconvolved cancer-specific gene expression by CMScaller, and we then compared the results from the same cell mixtures that were not subject to deconvolution (**Figure S1d**). We were able to accurately predict the proportion of cancer cell RNA in the cellRNA mixtures for all samples (**Figure S1d**). Therefore, mixtures containing as low as 0.1% cancer cell RNA could be stratified into the correct CMS after deconvolution (**Figure S1d**). An *in silico* mixing experiment was then performed to assess the accuracy of the CMS classifier on a larger deconvolved transcriptomic dataset. First, CRC tumor bulk RNA data from The Cancer Genome Atlas (TCGA) data were mixed *in silico* with bulk RNA-seq data derived from plasma EVs of healthy individuals. To generate such “synthetic mixtures”, the average read counts from normal plasma evRNA samples were computationally spiked in with RNA reads from CRC TCGA data using predefined vectors to obtain mixtures with different cancer portion ratios ranging from 0.01% to 10%. CIBERSORTx was used to predict the proportion of cancer-derived RNA (**Figure S2a**, P value<0.0001). The imputed proportions of each component (cancer and normal) is required as input for CIBERSORTx and CODEFACS to predict the sample-specific gene expression of cancer and normal groups. Next, the CMS classifier was applied to the deconvolved cancer specific expression, which showed that the assigned CMS subtype of the original TCGA sample could be successfully predicted following *in silico* deconvolution (**Figures S2b**). Synthetic mixture samples with lower proportions of cancer RNA were more likely to have discordant subtypes between the original sample and the deconvolved mixture (**Figure S2c**). Compared to CIBERSORTx, CODEFACS showed better performance than in predicting CMS4 whereas similar performance on CMS1-3 (**Figure S2b**). In addition, it lowered the proportion of discordant prediction of molecular subtypes to 1.2% compared to 2% for CIBERSORTx (**Figure S2c**).

**Figure 1.**
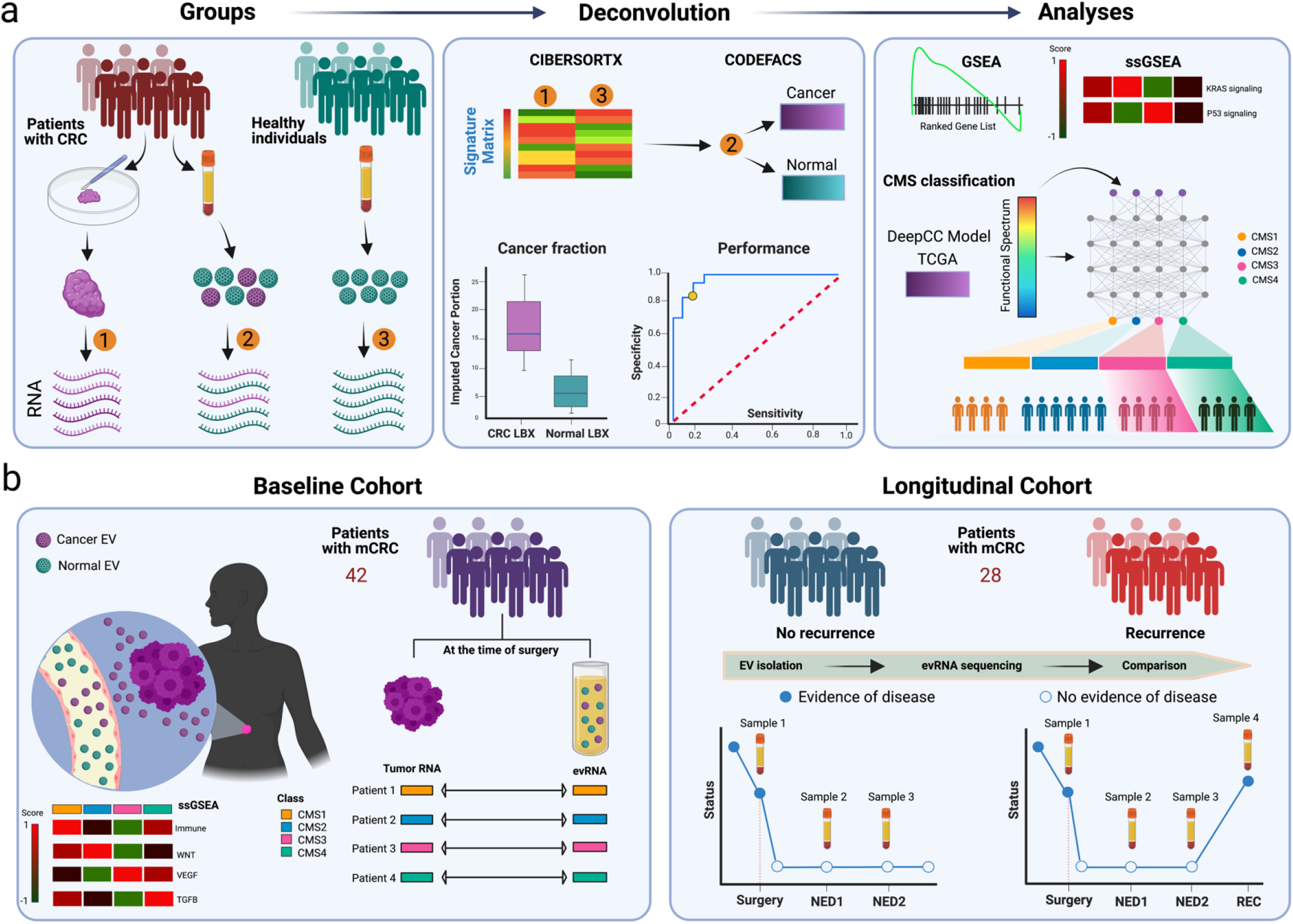
Performance of CMS subtyping of liquid biopsies from patients with mCRC. **a**, A visual overview of workflow. RNA-seq was performed on tumor samples and plasma EV of cancer patients as well as plasma EV of healthy controls. For deconvolution with CIBERSORTx and CODEFACS, a signature matrix was created using genes that are enriched in the tumor samples and in the plasma EV of healthy controls. Deconvolution could impute the proportion of cancer present in bulk plasma evRNA, which also allowed for generation of ROC curve based on whether cancer RNA was present in the sample. Deconvolved plasma EV profiles were also used in gene set enrichment analysis (GSEA) and artificial neural network to assign the CMS groups, which were then compared to the subtypes of the tumor samples. **b**, Cohorts of CRC patients analyzed in this study. In the baseline cohort, molecular subtypes of tumor tissues and their matched plasma evRNA samples were compared. In the longitudinal cohort, molecular subtype switch and emerging changes at the gene and pathway level were evaluated and compared, at each serial point, in patients with and without recurrence.

After establishing the pipeline based on *in vitro* and *in silico* studies, we aimed to analyze the feasibility of our approach in patient-derived liquid biopsies (**Figure 1a)**. We divided our samples into two CRC cohorts (**Figure 1b**). First, we utilized 42 baseline liquid biopsies from mCRC patients with paired tissue and 100 liquid biopsies from healthy individuals. Second, we applied our approach to longitudinal plasma evRNA samples from 28 patients, 12 with no recurrence and 16 with recurrence during the evaluated timeframe (**Figure 1b**). Samples from 12 patients with no recurrence, and 12 out of 16 with recurrence, were used for differential gene expression and gene set enrichment analyses. Longitudinal samples from 4 remaining patients with recurrence were used as examples for CMS switching.

In the baseline cohort, after deconvolution, the interquartile range of imputed proportions of cancer-derived RNA was 0.48 (median 1.52%). Receiver operating characteristic (ROC) curve for prediction of proportion of cancer-derived RNA showed an area under the curve (AUC) 0.961 with specificity of 0.96 and sensitivity of 0.9 at imputed cancer proportion threshold of 0.9% (**Figure S3a-b**). These results suggest that our EVs RNA deconvolution pipeline can accurately detect the presence of cancer even if the proportion of cancer-derived transcripts in the circulating evRNA transcriptome is as low as 1%.

### Plasma evRNA can be used to predict tumor transcriptomic subtype and track subtype evolution in longitudinal samples

In CRC, the tumor expression based CMS classification system has become widely accepted and has shown prognostic as well as predictive potential [25]. This classification system is based on tissue-derived transcriptome data. To test if the deconvolution pipeline can be used to predict tumor subtype through liquid biopsy, we applied our evRNA deconvolution pipeline on the aforementioned 42 plasma CRC plasma samples from mCRC patients with matching tissue (**Figure 2a-b**). The bulk evRNA without the deconvolution (mixture) was used as a control in CMS prediction (**Figure 2a**). We were able to correctly predict CMS of the tumor of origin in 71% (30/42) of cases. This concordance was significantly higher than the random concordance rate of 22%. Interestingly, the concordance between tumor tissue and liquid biopsy was as high as 84% (26/31) in patients with tumor purity higher than 10% (**Figure 2a**). Of note, most tissue samples with tumor purity less than 13% were classified as CMS4 (**Figure 2a**). Overall, in our evRNA cohort, 7.1% (3/42) of the patients’ tumors were classified as CMS1, 40.4% (17/42) as CMS2, 19% (8/42) as CMS3, and 33.3% (14/42) as CMS4 (**Figure 2a**). With assigning molecular subtypes of tumor tissues to their matched liquid biopsy samples, we showed that CMS specific pathways are differentially regulated in liquid biopsy samples (**Figure 2c-g**).

**Figure 2.**
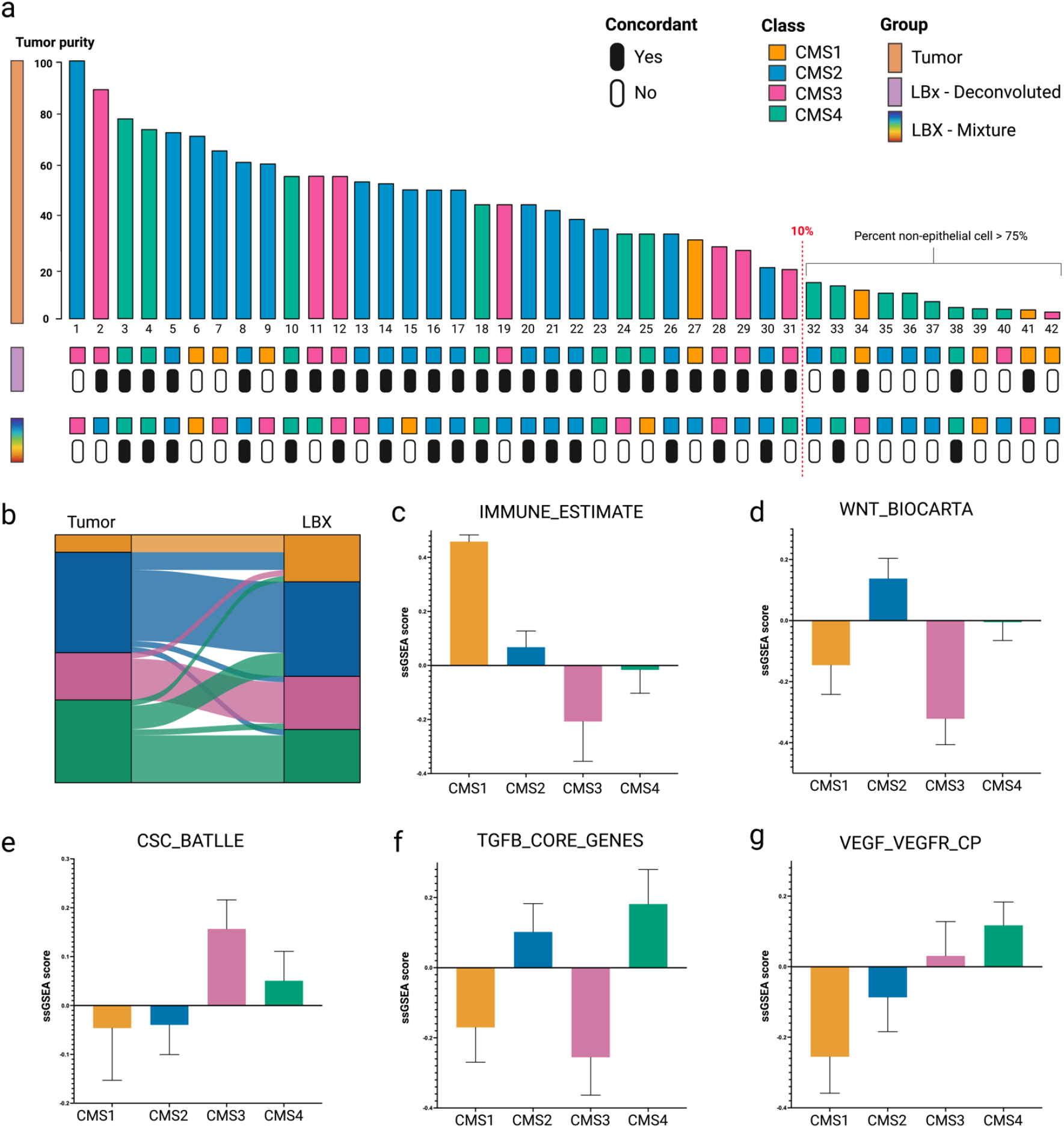
Application of RNA deconvolution in plasma from CRC patients. **a**, Summary of transcriptomic subtyping of liquid biopsy samples and their corresponding tumors. Each column represents a patient and columns are ordered by tumor purity. Top row represents the CMS group of each solid tumor, the middle row represents the prediction for deconvolved plasma EV, and the bottom row represents the prediction for bulk plasma EV. If the predicted subtype of the liquid biopsy is concordant with the tumor sample, the column is marked by black box. **b**, Sankey diagram depicting CMS classification for tumor (left) and paired liquid biopsy (right) using CODEFACS. **c-g**, single sample GSEA analysis for some of major pathways elevated at each CMS.

We then applied our evRNA deconvolution pipeline to 15 longitudinal samples collected from four mCRC patients from the recurrence cohort (**Figure 3**). In patient ID 1, an evRNA liquid biopsy performed after neoadjuvant treatment for a single metachronous metastasis assigned the tumor to subtype CMS2. After liver resection, both CT scans and cfDNA liquid biopsies were negative. However, evRNA liquid biopsy was positive 3 months before radiological manifestation of disease recurrence and our algorithm initially assigned the recurrence to CMS2. At the time of radiological progression, however, a switch to CMS4 was observed, while cfDNA liquid biopsies continued to be negative. (**Figure 3a**). A similar switch from CMS2 to CMS4 was detected at the time of radiological progression in patient ID 2. Interestingly, the CMS switch in this patient was associated with the identification of mutant *KRAS* in cfDNA. (**Figure 3b**). Patient ID 3 presented with multiple synchronous liver metastases. After neoadjuvant therapy and partial metastasectomy combined with radiofrequency ablation, evRNA liquid biopsy initially assigned a CMS2. The subsequent clinical course continued to demonstrate a relatively stable CMS2 subtype on repeated sampling. Nonetheless, a switch to CMS 1 was observed only a few weeks preceding a CT scan that demonstrated a substantial progression in the liver. (**Figure 3c**). A similar pattern was observed in patient ID 4 who was initially classified as CMS2, on two consecutive liquid biopsies after neoadjuvant treatment and partial hepatectomy. However, he later switched into CMS1 three months prior to detection of radiological progression in the liver and maintained as CMS1 on a consecutive draw. After microwave ablation, the patient had a subsequent draw positive for mutant *KRAS* in cfDNA which was followed by a second relapse in the liver and a switch into CMS4 (**Figure 3d**). Thus, in all four cases, we observed a “subtype switch” in evRNA CMS assignment which predates the appearance of therapeutic resistance and radiological progression.

**Figure 3.**
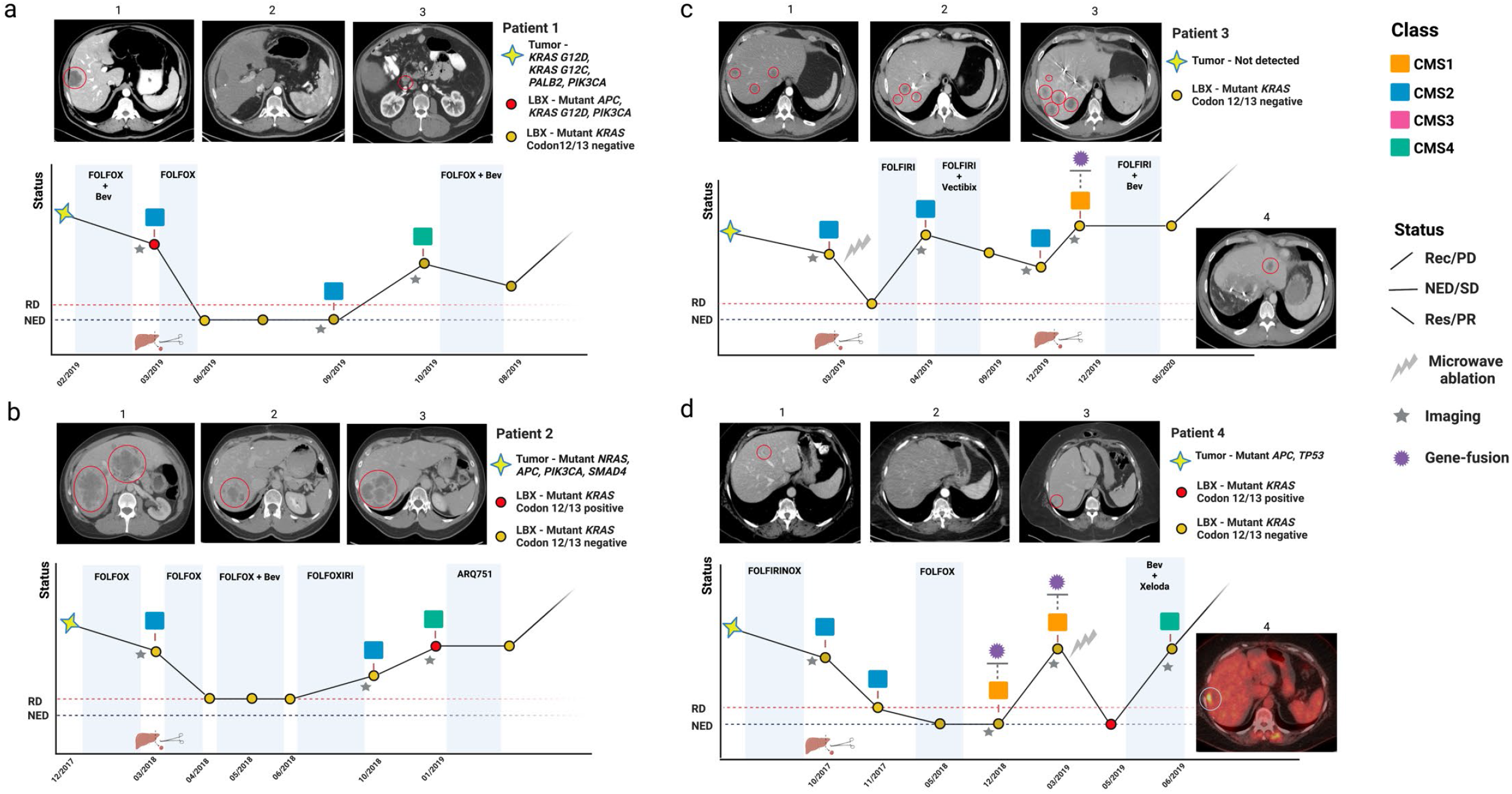
Longitudinal monitoring of CMS changes in mCRC patients. Timeline showing disease history of four mCRC patients. (**a** and **b**) In patients one and two, EVs-based CMS was determined at baseline, before and at the time of radiological progression. **c**, patient three, EVs-based CMS was determined at baseline and at five additional time points. Disease responses according to RECIST 1.1 [45] criteria are shown in the y axis. **d**, in patient four, EVs-based CMS was predicted at baseline and in four additional longitudinal draws. An upward segment between two time points represents recurrence/progressive disease (Rec/PD), a flat segment represents stable disease (SD) or non-evidence of disease (NED) and a downward segment represents complete response/partial response (CR/PR).

In addition to CMS classification, we investigated whether our approach was able to capture transcriptomic changes, at the gene and pathway levels which could help elucidate molecular underpinnings of disease relapse. For this analysis, we used our longitudinal cohort consisting of 24 metastatic patients that underwent hepatectomy. 12 patients showed no evidence of disease (NED) within a median follow up of 425 days (range 175-1578 days) from the time of hepatectomy while the other 12 patients recurred within a median follow up of 435 days after surgery (range 97 to 1132 days). For patients with no recurrence, two serial draws were assessed in the post-operative period, both at the time of clinical NED (**Figure 4a**). For patients with recurrence, two follow-up draws during the NED interval, and a third on clinical recurrence were analyzed (**Figure 4a**). Proportions of cancer-derived evRNA were analyzed at each time point using our deconvolution pipeline. These proportions were significantly higher at recurrence compared to previous draws and were lower in NED draw 2 (NED2) for patients without recurrence (**Figure 4a-b**). We then performed Gene Set Enrichment Analysis (GSEA) at each longitudinal draw and compared the average pathway scores across all 12 patients from each group within deconvoluted evRNA. Multiple pathways showed opposing trends between these two groups, including KRAS signaling and interferon gamma response, amongst others (**Figure 4a-b**). Patients that underwent clinical relapse had an upregulation of both pathways at recurrence, while patients that remained NED, showed a decrease from draw 1 to draw 2. Additional differentially enriched pathways are illustrated in **Figure S4**. Because of the opposite trends observed in NED2 by pathway analysis, we examined differences in gene expression at this draw, for both groups. Several cancer-associated genes were differentially expressed in patients with recurrence, including transcripts corresponding to the oncogenes *MET, MYC* and *KRAS* (**Figure 4c**).

**Figure 4.**
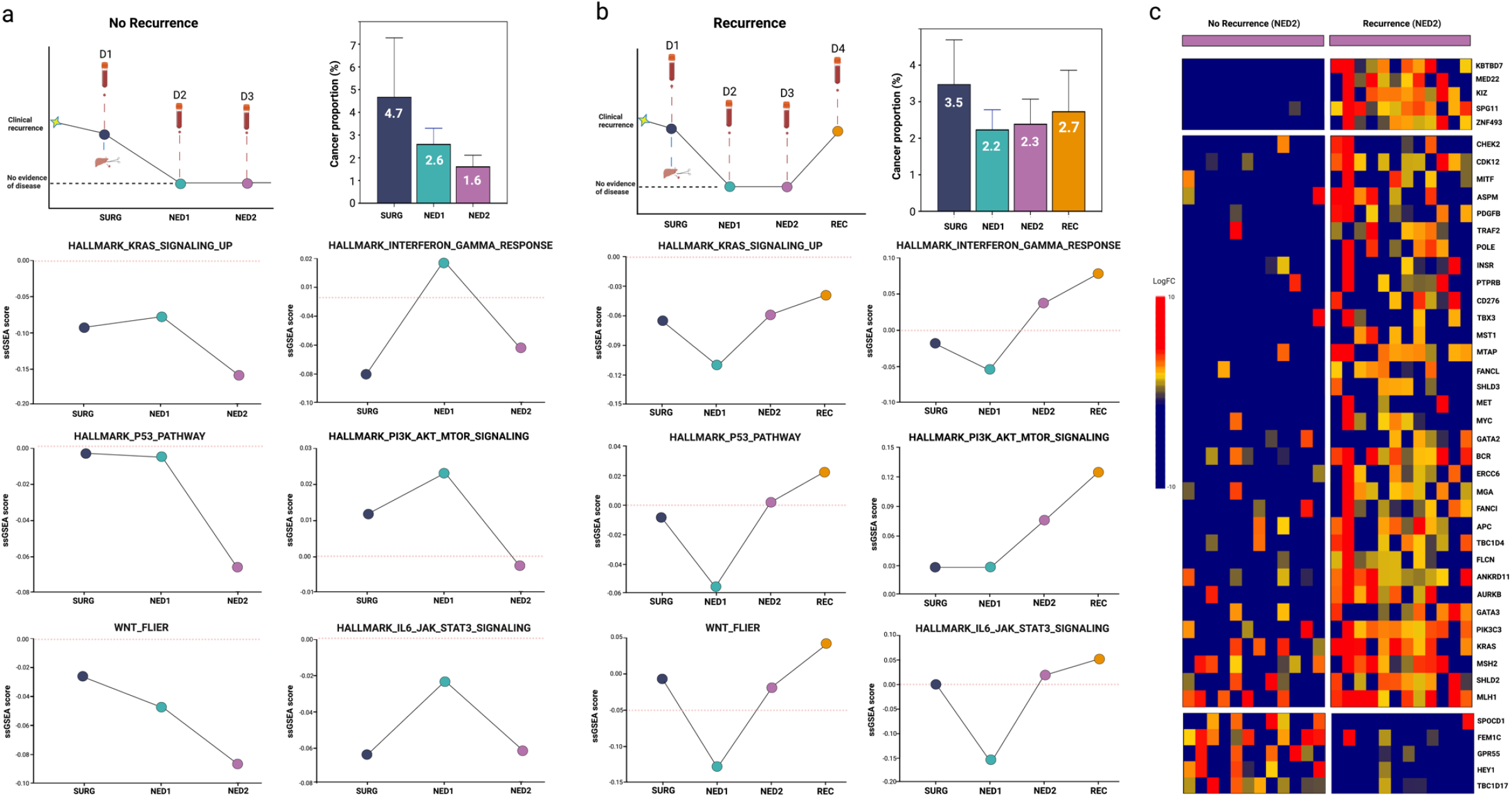
Longitudinal monitoring of genes and pathways changes in mCRC patients. Representation of longitudinal timeline for patients with no recurrence (**a**) and recurrence (**b**). Average cancer proportions for all patients at each time point is shown at the right of the timeline and average of single sample gene set enrichment analysis (ssGSEA) scores for all cases is shown below. **c**, Differential gene expression at NED2 for patients with no recurrence (yellow) and recurrence (red). Patients are shown as columns.

### evRNA can identify expressed gene fusions and neoepitopes

Chromosomal rearrangements have been well documented in colorectal cancer and can lead to the formation of gene fusions, offering new targets for personalized therapy [26, 27]. We evaluated the present of gene fusions in evRNA from our baseline and longitudinal cohorts by using three bioinformatic pipelines (Arriba, Pizzly pair and Fusioncatcher, see Methods). In our baseline cohort, three gene fusions, *DOCK4:IMMP2L, SLC6A6:XPC* and *TFG:ADGRG7*, were detected independently in three samples, both in evRNA and in matched tumor tissues (**Figure S5a-c**). The gene fusion *DOCK4::IMMP2L* is of particular interest, since *IMMP2L* somatic breakpoints have been reported in inflammatory bowel disease-associated CRC, and somatic rearrangements have been also described for this gene [28, 29]. The gene fusion *GATC::COX6A1* could be identified in two independent evRNA samples but not in the matched tissues. This could be ascribable to the limited cancer region sampled for sequencing (**Figure S5d**). In our longitudinal cohort, we identified 39 total gene fusions in 25 out of 84 samples (30%) which were called by at least two of the algorithms. Interestingly, in patients with no recurrence, no gene fusions were detected in NED2 evRNA samples, while in patients with recurrence, fusions were present in NED2 evRNA and at recurrence (**Figure 5a, Table S1**). Additionally, six gene fusions were detected in MAPK-related genes, five in *ITGB3* and four in *CHIC2*. Interestingly, *VCPIP1:MAPK1, PTPRK:FAM120B* and *KRAS:SENP6* were identified at in evRNA samples of patient 7 at baseline, NED2 and at recurrence, respectively (**Figure 5b-d)**.

**Figure 5.**
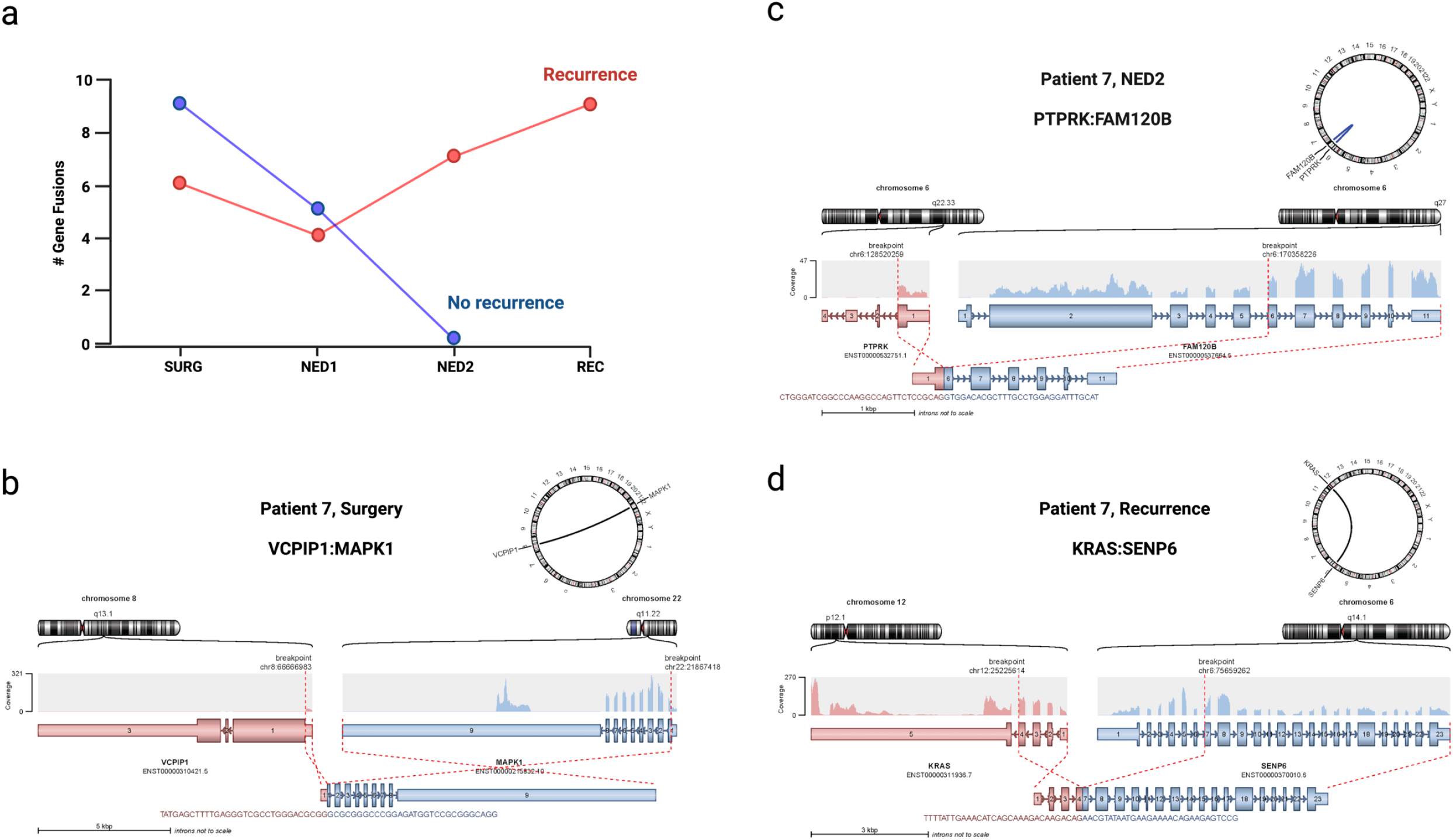
Prediction of gene fusions in plasma derived EV. **a**, quantification of gene fusions in patients with and without recurrence. **b-d**, Structural representation of gene fusions and circle plot depicting main chromosomal aberrations in a patient with recurrence (patient IDA 7). Fusions at baseline (**b**, VCPIP1:MAPK1), NED2 (**c**, PTPRK;FAM120B) and recurrence (**d**, KRAS:SENP6) are depicted.

Furthermore, we investigated the presence of neoepitopes from gene fusions, insertions/deletions (INDELs) and exitrons in our initial cohort of 42 baseline samples while comparing liquid biopsy calls to paired tissue samples. Through the pipeline NeoFuse we were able to predict the peptide HQDQAVSL from the gene fusion GATC:COX6A1 (**Table S2**). ScanNeo was utilized to predict INDELs-derived neoepitopes. With this pipeline we predicted matched plasma and tissue neoepitopes in 15/42 patients, and a total of 466 paired neoepitopes with a <500mM mutant binding affinity were found. Among these, 181 neoepitopes from 9 patients were predicted to have <100nM binding affinity and 158 from 10 patients showed a <50nM binding (**Figure S3c, Table S2**). We then applied ScanExitron, a computational pipeline to identify exitron spicing events. We were able to identify seven exitrons in both plasma and matched tumor samples. Among these, two exitrons contained one and two predicted neoantigens, respectively (**Table S2**).

## Discussion

In this study, we coupled low-input RNA-seq data obtained from circulating EVs with a bioinformatic deconvolution pipeline and deep learning algorithms in order to interrogate multiple tumor-specific transcriptomic features *sans* the requirement for tissue biopsies. Shi and colleagues are credited with the first demonstration that transcriptomic analyses of circulating EVs can be used to predict the response of advanced melanomas to immune checkpoint inhibitor (ICI) immunotherapy [30]. Notably, response to ICIs was associated with an enrichment for transcripts related to immune pathways within post-treatment EV samples, and the latter were principally contributed by non-tumor sources (i.e., immune cells). Although this prior study principally focused on the non-tumor EV transcriptome, it laid the foundational basis for exploring the possibility that evRNAs might represent a rich resource for mining cancer-specific information, so long as the cancer-specific compartment could be reliably extracted from the composite pool of circulating EVs.

Our study demonstrates that bioinformatic deconvolution of cancer-specific RNAs in plasma EVs is indeed feasible, to as low a fraction as only 1% of cancer-derived transcripts in the circulating evRNA transcriptome. While the current limit of detection in our algorithm is at 1% of cancer-derived transcripts within the evRNA pool, ongoing improvements in the pipeline is likely to further reduce this threshold, enabling us to track patients in the minimal residual disease (MRD) setting. Using our deconvolution pipeline, we were able to successfully predict tumor molecular subtypes in the majority of CRC cases analyzed, using tissue-based RNA profiling as the “ground truth”. The discrepancies identified between blood and tissue in the case of CRC are likely the combined result of underlying heterogeneity, where region-specific sampling of the tissue impacts the predicted subtype, and the impact of non-tumor components (stroma and immune cells) to the subtype annotation. Importantly, our pipeline used evRNA signatures not only for classifying tumors into established molecular subtypes at baseline, but also for longitudinal monitoring of changes in aforementioned molecular subtypes over time. For example, longitudinal monitoring of evRNAs in a cohort of CRC patients was able to identify changes in molecular subtype prior to onset of imaging-based progression, underscoring the importance of this approach to act as a sentinel for emergence of treatment resistance. While the association between tumor subtype switching and treatment resistance is well recognized [13], such monitoring has mostly been restricted to serial tissue samples, which is neither universally feasible nor cost effective. Liquid biopsies may facilitate the ability to detect evolution in CMS subtypes without the need of invasive biopsies. For example, a switch from CMS2 to CMS1 was detected before radiological progression in cases 3 and 4, thus opening new possibilities for potential immune therapy intervention for these patients, as CMS1 stratification has been reported to be more responsive to immune therapies [31].

Our approach was not only able to detect changes in CMS but also gene and pathway level changes over time. When comparing patients that progressed with those that did not during the time monitored, we were able to detect subtle differences in blood that could indicate residual disease. For instance, *KRAS, MYC*, and *MET*, were overexpressed at NED2 in patients with recurrence. *MET* amplification has been associated with resistance to anti-EGFR therapy in CRC [32]. Similarly, elevated expression of *MYC* and *KRAS* have been related to resistance to EGFR therapy in mCRC patients [33, 34]. Interestingly, patients with recurrence showed increase on various genes involved in DNA damage repair which may suggest an unstable genome.

Finally, the identification of gene fusions and neoepitopes in blood brings a unique opportunity for precision oncology for advanced cancer patients. In CRCs, expressed fusions are often actionable [35-37], but their identification can be limited in tissue-based assays. A blood-based assay provides redundant opportunities for assessing fusions, including over the course of time. In addition, our results indicate that the detection of gene fusions could be used as a potential biomarker for recurrence and tumor monitoring in mCRC patients. Moreover, the identification of potential neoepitopes without the prerequisite for tissue biopsies is of considerable translational relevance in patients receiving neoantigen targeted therapies (such as vaccines or engineered T cell receptors), where both *de novo* epitopes and newly emerged candidates can be tracked longitudinally over time and the therapeutic modulated accordingly.

Our data underpins the utility of plasma evRNA as a multifaceted liquid biopsy analyte in cancer patients, complementing the well-established of DNA-based profiling. Transcriptomic subtyping of tumors through liquid biopsy can be of particular interest when routine tissue biopsy is not feasible. This approach can be used to predict molecular subtypes of solid tumors at baseline, but more importantly, serve as a platform for identifying determinants of response and resistance, as well as potentially anticipate clinical recurrence in serial samples. In our companion study by Van Morris et al, we demonstrate the utility of our evRNA based profiling and deconvolution pipeline for identifying correlates of response to a novel “triple therapy” regimen in *BRAF*^V600E,^ mutant colorectal cancer. The two studies together lay the foundational basis for extrapolating the power of liquid biopsy-based monitoring of cancer patients using direct RNA profiling.

## Methods

### Cell lines and culturing

Ten established CRC cell lines, LOVO (CMS1), COLO201 (CMS1), LS1034 (CMS2), SNUC1 (CMS2), HT29 (CMS3), LS513 (CMS3), WiDr (CMS3), HCT116 (CMS4), LS123 (CMS4), SW480 (CMS4) from primary or metastatic CRC tissue were used in this study. A CRC neoplastic cell line, HCC2998, with mixed molecular subtypes and 481CoN, a normal colon cell line, were used as controls. All cell lines were maintained either in RPMI-1640 or DMEM medium with 10% FBS, 100 units/mL penicillin and 100 µg/mL streptomycin (Gibco, Life Technologies, Carlsbad, CA, USA) at 37°C and 5% CO2 in a humidified incubator. Before EV isolation, cell lines were cultured in HYPERflasks (Corning Inc, Corning, NY, USA) in medium with 10% exosome-free FBS for at least 48 hours and conditioned media was then harvested. Next, 500 mL of media per cell line was centrifuged serially at 1000 rpm for 10 minutes and 5000 rpm for 5 minutes at 4°C to remove all cellular debris. The media was then passed through a 400-nm filter to remove any remaining debris and transferred to 50-mL tubes. The media was then ultra-centrifuged at 36,000 rpm at 4°C overnight. The resulting pellet was resuspended in PBS with a subsequent ultra-centrifugation for 2-4 hours at 36,000 rpm. The EV pellet was resuspended in 600 uL PBS.

### Patient samples

One hundred and forty-two plasma samples from 70 patients (42 for the baseline cohort and 100 for the longitudinal cohort) with histologically confirmed CRC who were diagnosed and treated at the University of Texas MD Anderson Cancer Center (UTMDACC) between 8/8/2014 and 8/29/2019 were enrolled for this study. Informed consent was obtained following protocol LAB10-0982 in accordance with standard ethical guidelines approved by the institutional review board (IRB) and the Declaration of Helsinki. Blood samples were collected preoperatively on the day of surgery, tissue samples were collected at the time of surgical resection. Blood samples from one hundred healthy individuals were collected (matched based on age/gender).

### Next-generation sequencing library construction for tumor tissues

Following RNA extraction, all tissue samples were processed using the KAPA Stranded RNA-Seq kit with RiboErase (HMR) (KAPA, Roche, cat. # 08098140702). Libraries were constructed according to the manufacturer protocol and were then quantified using an Agilent 2100 Tapestation. Libraries were then sequenced using 150-bp paired-end runs on a NextSeq 500 sequencer (Illumina, San Diego, CA, USA).

### EV Isolation from human plasma samples

Three 8ml EDTA Vacutainer tubes (BD) of blood were collected from each case. Plasma was separated by centrifuging blood samples at 2500 rpm for 10 minutes at room temperature (RT), followed by an additional 5-minute 5000 rpm centrifugation at 4°C to remove cellular debris. The plasma was ultracentrifugated overnight at 36,000 rpm. The EV pellet was then washed with PBS and centrifuged at 36,000 rpm for 2-4 hours. The supernatant was then discarded, and the EV pellets were resuspended in 600ul PBS for long term storage at -80°C.

### RNA extraction and next generation sequencing

200 μL of resuspended EV were used for RNA extraction using the total exosome RNA & Protein Isolation kit (Invitrogen, cat. # 4478545) following the manufacturer’s protocol. Small RNA species such as miRNAs were removed during the washing steps. Contaminating DNA was removed using DNase I (Invitrogen, cat. #18047019). cDNA amplification was performed using the SMART-Seq® v4 Ultra® low input RNA kit (Takara, cat. # 634891) with sixteen cycles of PCR. cDNA samples were then diluted to 0.2-0.3 ng per uL and 1 ng of cDNA was carried over into library preparation using the Nextera XT DNA Library Preparation Kit (Illumina, cat. # FC-131-1096) following the manufacturer’s protocol. Libraries were quantified with an Agilent 2100 Tapestation and sequenced in a NextSeq 500 sequencer. Samples with fewer than 1 million coding reads and high percentage of globin RNA (gRNA) and ribosomal RNA (rRNA), library prep was repeated with an additional globin RNA and ribosomal RNA depletion step using riboPOOLs gRNA/rRNA depletion kit (siTOOLsBiotech, cat. # dp-K096-001002).

### Creation of custom gene set for CMS classification of CRC cell line EV

The use of the CMS gene set provided in the ‘CMScaller’ package [18] led to inaccurate classification in two of eleven cell line EV (data not shown). In order to improve the performance of nearest template prediction for EV, the preexisting gene list was subset by taking the genes that are highly expressed in the cells and EV of each CMS group. For cell lines and EV of each CMS group, differentially expressed genes compared to cell lines and EV of other CMS groups were obtained using the ‘limma’ package. Only those genes in the ‘CMScaller’ gene set that were upregulated in cell lines and EV of each CMS group with fold change of 2 or greater and adjusted *P*-value < 0.05 were included in the new gene set.

### Artificial mixing experiments

Synthetic mixtures from CRC TCGA RNA sequencing data were generated for performance assessment of deconvolution methods. We first evaluated the correlation between the observed and expected cancer proportions. We then assessed the correlation between observed cancer proportions and molecular subtypes concordance between liquid biopsy and tissue samples. To generate synthetic mixture, the average read counts from normal plasma evRNA samples were computationally spiked in with RNA reads from CRC TCGA data using predefined vectors to obtain mixtures with different cancer portion ratios ranging from 0.01% to 10%. Multiple linear regression was used to assess the linear relationship between the observed and expected cancer levels. We then assigned each deconvolved expression profile to a molecular subtype by using pure TCGA expression data as the reference for CMScaller.

### RNA-Seq data analysis, deconvolution and molecular classification

QC was performed on FASTQ files using MultiQC. Raw reads were aligned to the human genome reference (hg19) using STAR. Deep cancer subtype classification (DeepCC) [21] was used to classify the transcriptomic profiles of tumor tissues. In summary, we trained a DeepCC classifier using a TCGA CRC data set (*n*□=□456) and a multilayer artificial neural network (ANN) as previously described by Gao et al [21]. We then obtained the functional spectra from tumor expression profiles and used them as input to predict molecular subtypes for tumor samples.

To predict the molecular subtypes from liquid biopsy samples, we first bioinformatically deconvolved the bulk gene expression profiles in order to exclude the transcriptomic profile of normal EV. We applied two deconvolution methods CIBERSORTx [22] and CODEFACS [24] to the normalized reads to impute proportions of cancer in the plasma of CRC patients and to assign molecular subtypes to the gene expression profile of the cancer-specific compartment. Both deconvolution methods were tested, by first *in silico* experiments and then deconvoluting cellular RNA mixtures with known proportions (**Figure S1 and Figure S2**). CMS subtyping was performed by two different classifiers, one CMScaller (R package CMScaller) which uses CMS genes predicted by the nearest template prediction method and another by ANN-based classification [19, 21]. For the latter, gene expression profiles of CRC-stratified cell lines were used to train a DeepCC classifier (**Figure S1**).

Next, we performed deconvolution of bulk RNA isolated from the plasma EV of CRC patients (**Figure 1**). Deconvolution was first used for the assessment of cancer evRNA abundance (**Figure S3**). To evaluate and compare cancer evRNA levels between CRC and normal plasma samples, samples from healthy individuals from the first cohort were randomly divided into two groups. A group of 29 normal samples was used as the predefined normal group for both deconvolution methods. For the accurate assessment of cancer evRNA levels, 29 CRC plasma samples were selected (based on sample quality) to compare with the remaining normal samples (29 samples, control group) (**Figures S3**).

We then used the cancer-specific compartment of the deconvolved transcriptomic profiles to predict the CMS class by ANN-based classification. CRC TCGA RNA sequencing data was used to train a classifier as described for the tumor tissue CMS classification (**Figures 1**). For the deconvolution by CODEFACS, a signature matrix, which is comprised of genes that are enriched in each group of interest, expression profiles of TCGA RNA sequencing data and 100 normal plasma EV were used (**Figure 1a**). High-resolution expression deconvolution was then applied to 42 EV samples from the same patients with tumor samples (**Figure 1a**). Paired Student’s *t* test was used to calculate *P*-values when comparing imputed cancer portions.

### Gene fusions and neoantigens prediction

Gene fusions were analyzed from 42 tissue samples, matched liquid biopsies and 58 healthy controls for our first analysis and additional 142 liquid biopsies (100 mCRC and 42 healthy) from our second phase of the study. Three different methods were used to detect gene fusions and settings criteria was kept as: a) Arriba [38], confidence of medium or high, discordant_mates ≥1, split reads_reads ≥2, b) Pizzly [39], paircount ≥1 and split count ≥2 and c) FusionCatcher [40], spanning_pairs ≥1 and spanning_unique_reads ≥2. Gene fusions present in healthy controls were removed from the analysis. Neoantigens were detected by NeoFuse [41] and only cases with binding affinity < 100nM were reported (**Figure S3c** and **Table S1**).

### Neoepitopes derived from INDELs and Exitrons

INDELs and exitrons were analyzed from 42 tissue samples, matched liquid biopsies and 58 healthy controls. INDELs were called by TransIndel [42] followed by neoepitopes analysis by ScanNeo [43] and calls present in healthy controls were removed. Exitrons were called by ScanExitron [44] and ScanNeo further predicted neoantigens (**Table S1**).

## Figure Legends

**Figure S1**. CRC cell lines EV transcriptome from Consensus molecular subtypes of cell lines and their EV are concordant. a, Heatmap representing the expression of CMS genes in cRNA and evRNA from 10 different colorectal cancer cell lines. Rows are ordered by genes in each CMS class and columns are ordered by CMS subtype and the P-value of nearest template prediction from CMScaller (white bar). b, Heatmap representation of CMS group of each cell line and its EV, with top row representing previously published subtypes. c, Scatter plot of actual proportion of cancer cellRNA against imputed proportion after deconvolution using CIBERSORTx. d, Heatmap representation of CMS group of each cellRNA mixture for cell lines of known molecular subtype before and after deconvolution and different subtyping methods.

**Figure S2**. Performance of deconvolution methods on artificial mixing experiments using TCGA CRC gene expression data. a, Correlation plot of actual proportion of cancer RNA and imputed cancer RNA using CIBERSORTx. b, Sankey diagram demonstrating the CMS group of original cancer samples on the left and the CMS group of deconvolved expression data from artificial mixtures (right) using CIBERSORTx (left) and CODEFACS (right). c, Violin plots of imputed cancer proportion in deconvolved expression data comparing discordant and concordant samples using CIBERSORTx (left) and CODEFACS (right).

**Figure S3**. Analysis of cancer proportions. a, Imputed cancer portions for CRC samples compared to normal evRNA. b, Receiver Operating Characteristic (ROC) curve analysis for the imputed cancer portions was performed in order to estimate the area under the curve (AUC) for CIBERSORTx. c, predicted neoepitopes identified in paired plasma and tissue samples and grouped by binding affinity.

**Figure S4. Application of RNA** deconvolution **in plasma from CRC patients**.

Representation of longitudinal timeline for patients with no recurrence (**a**) and recurrence (**b**). Differential GSEA analysis (GSEA Hallmark) at each longitudinal point is shown below. Each data point represents the average of all samples.

**Figure S5. Prediction of gene fusions in plasma derived EV**. Plasma, paired tumor and healthy controls were analyzed for gene fusions using Arriba, Pizzly and FusionCatcher. Structural representation of gene fusion and circle plot depicting main chromosomal aberrations. **a**, DOCK4:IMMP2L fusion was detected in patient ID 21 and supported by arriba: pair=4, reads=4, pizzly:pair=15, reads=11, and fusioncatcher: pair=7, reads=6. **b**, SLC6A6:XPC and antisense SLC6A6:XPC-AS1 fusions were detected in patient ID 24 and supported by arriba: pair=13,reads=24 and fusioncatcher: pair=16,reads=7 respectively. **c**, TFG:ADGRG7 fusion was detected in patient ID 27 and was supported by fusioncatcher: pair=2, reads=4 and pizzly: pair=1, reads=3. **d**, GATC:COX6A1 was detected in patients IDs 11 and 40 and was supported by arriba: pair =3, reads =2 and arriba: pair=7, reads =26, respectively.

**Table S1**. Predicted matched plasma and tissue neoantigens identified in gene fusions, INDELs and exitrons with a mutant binding affinity <100nM.

**Table S2**. Predicted gene fusions detected in mCRC patients with and without recurrence.

## Acknowledgments

We thank our patients and their families.

