## Supplementary Figures for "Transcriptomic Profiling of Plasma Extracellular Vesicles Enables Reliable Annotation of the Cancer-specific Transcriptome and Molecular Subtype"

### Figure S1

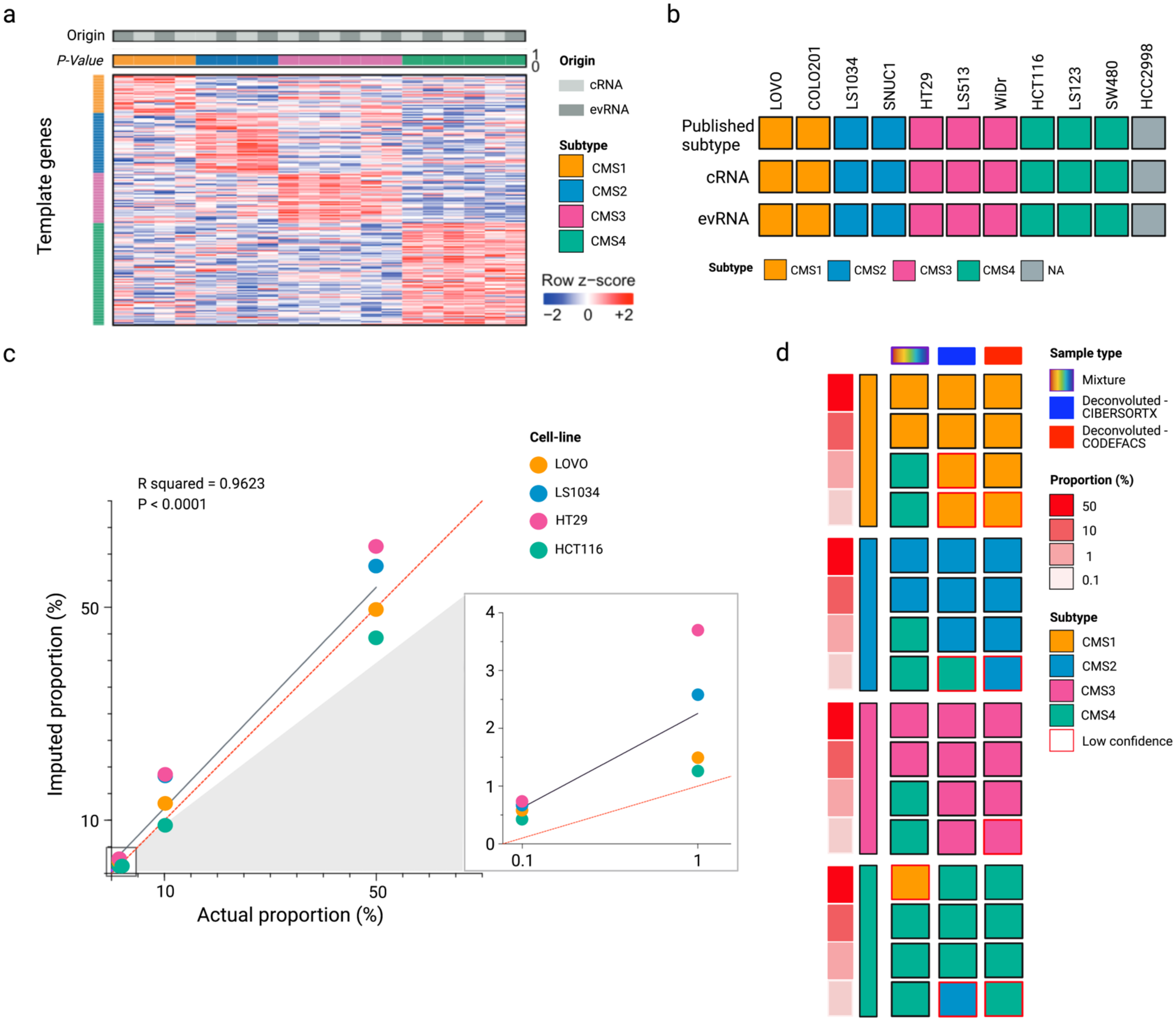


**Figure S1. CRC cell lines EV transcriptome from Consensus molecular subtypes of cell lines and their EV are concordant. a,** Heatmap representing the expression of CMS genes in cRNA and evRNA from 10 different colorectal cancer cell lines. Rows are ordered by genes in each CMS class and columns are ordered by CMS subtype and the P-value of nearest template prediction from CMScaller (white bar). **b,** Heatmap representation of CMS group of each cell line and its EV, with top row representing previously published subtypes. **c,** Scatter plot of actual proportion of cancer cellRNA against imputed proportion after deconvolution using CIBERSORTx. **d,** Heatmap representation of CMS group of each cellRNA mixture for cell lines of known molecular subtype before and after deconvolution and different subtyping methods.

### Figure S2

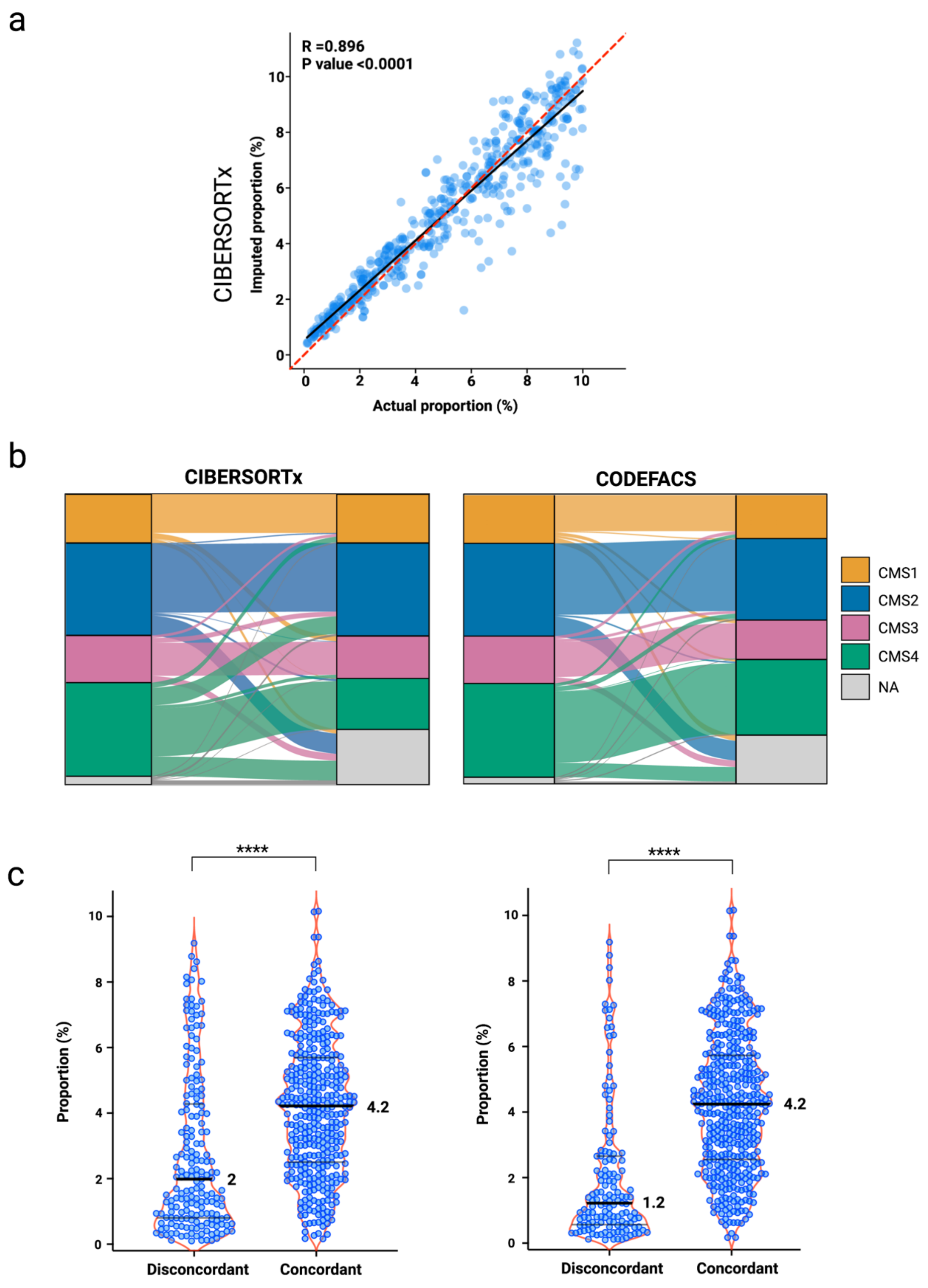


**Figure S2. Performance of deconvolution methods on artificial mixing experiments using TCGA CRC gene expression data. a,** Correlation plot of actual proportion of cancer RNA and imputed cancer RNA using CIBERSORTx. **b,** Sankey diagram demonstrating the CMS group of original cancer samples on the left and the CMS group of deconvolved expression data from artificial mixtures (right) using CIBERSORTx (left) and CODEFACS (right). **c,** Violin plots of imputed cancer proportion in deconvolved expression data comparing discordant and concordant samples using CIBERSORTx (left) and CODEFACS (right).

**Figure S3**

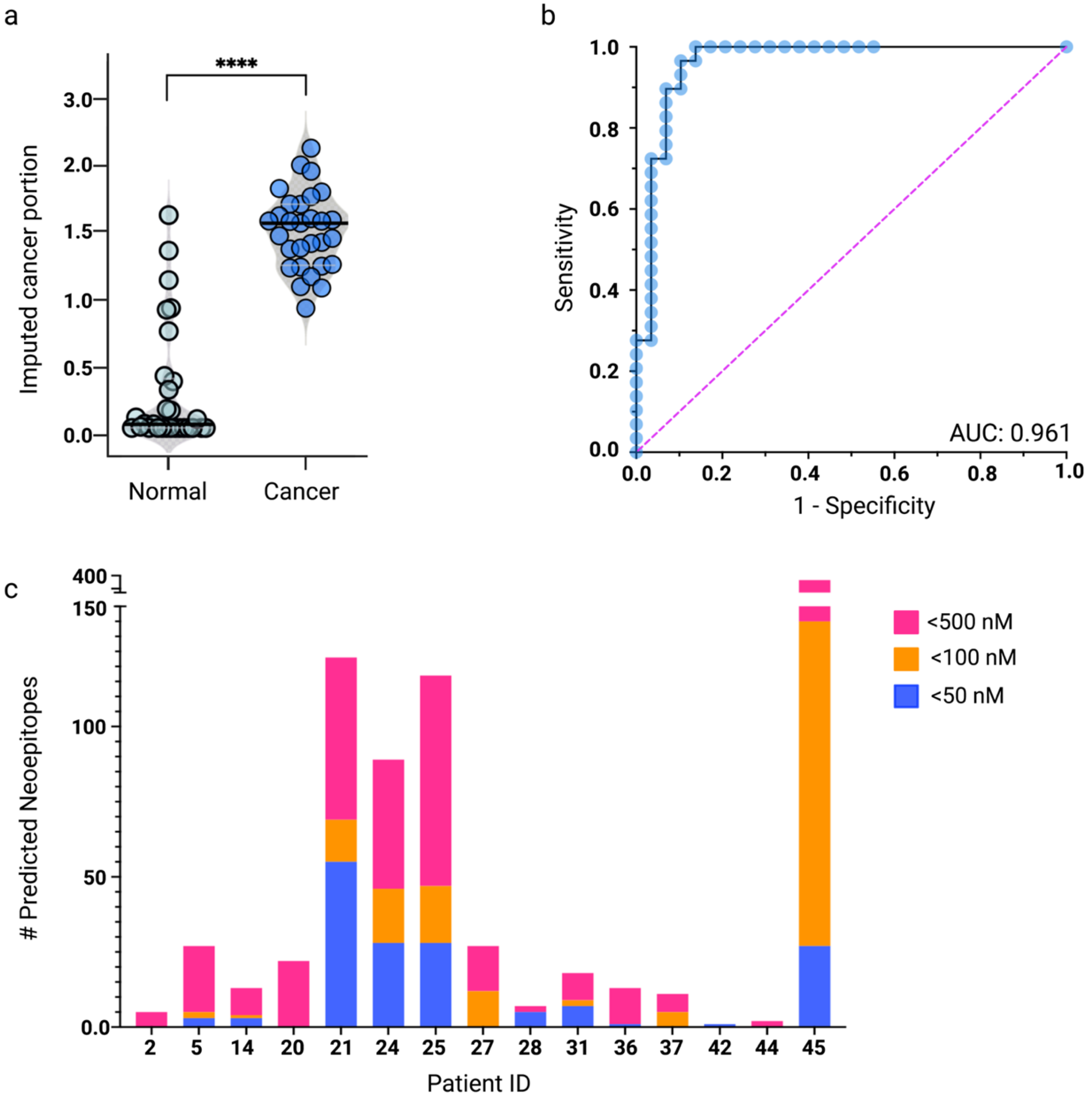


**Figure S3. Analysis of cancer proportions**. **a,** Imputed cancer portions for CRC samples compared to normal evRNA. **b,** Receiver Operating Characteristic (ROC) curve analysis for the imputed cancer portions was performed in order to estimate the area under the curve (AUC) for CIBERSORTx. **c,** predicted neoepitopes identified in paired plasma and tissue samples and grouped by binding affinity.

### Figure S4

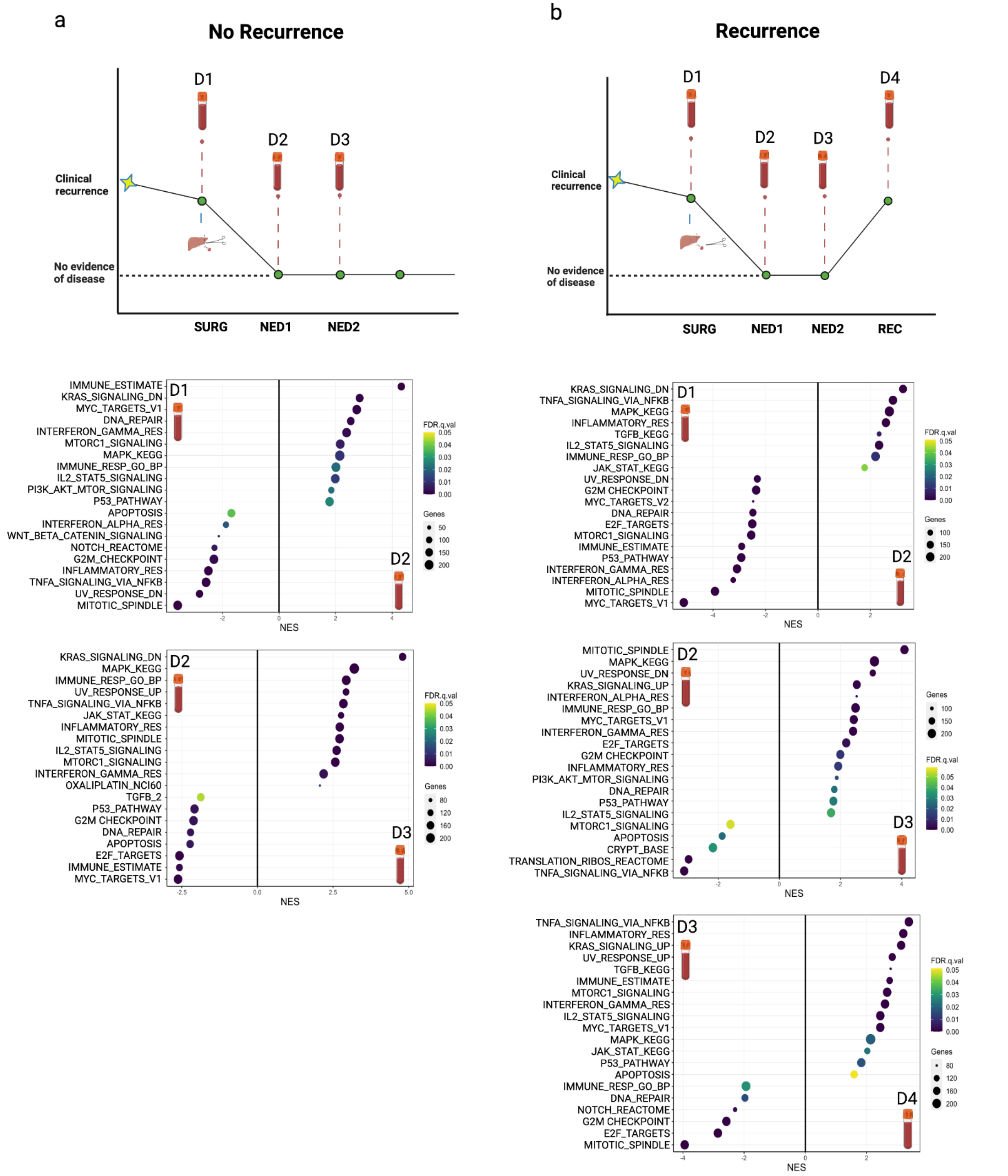


**Figure S4. Application of RNA deconvolution in plasma from PDAC and CRC patients.** Representation of longitudinal timeline for patients with no recurrence (**a**) and recurrence (**b**). Differential GSEA analysis (GSEA Hallmark) at each longitudinal point is shown below. Each data point represents the average of all samples.

### Figure S5

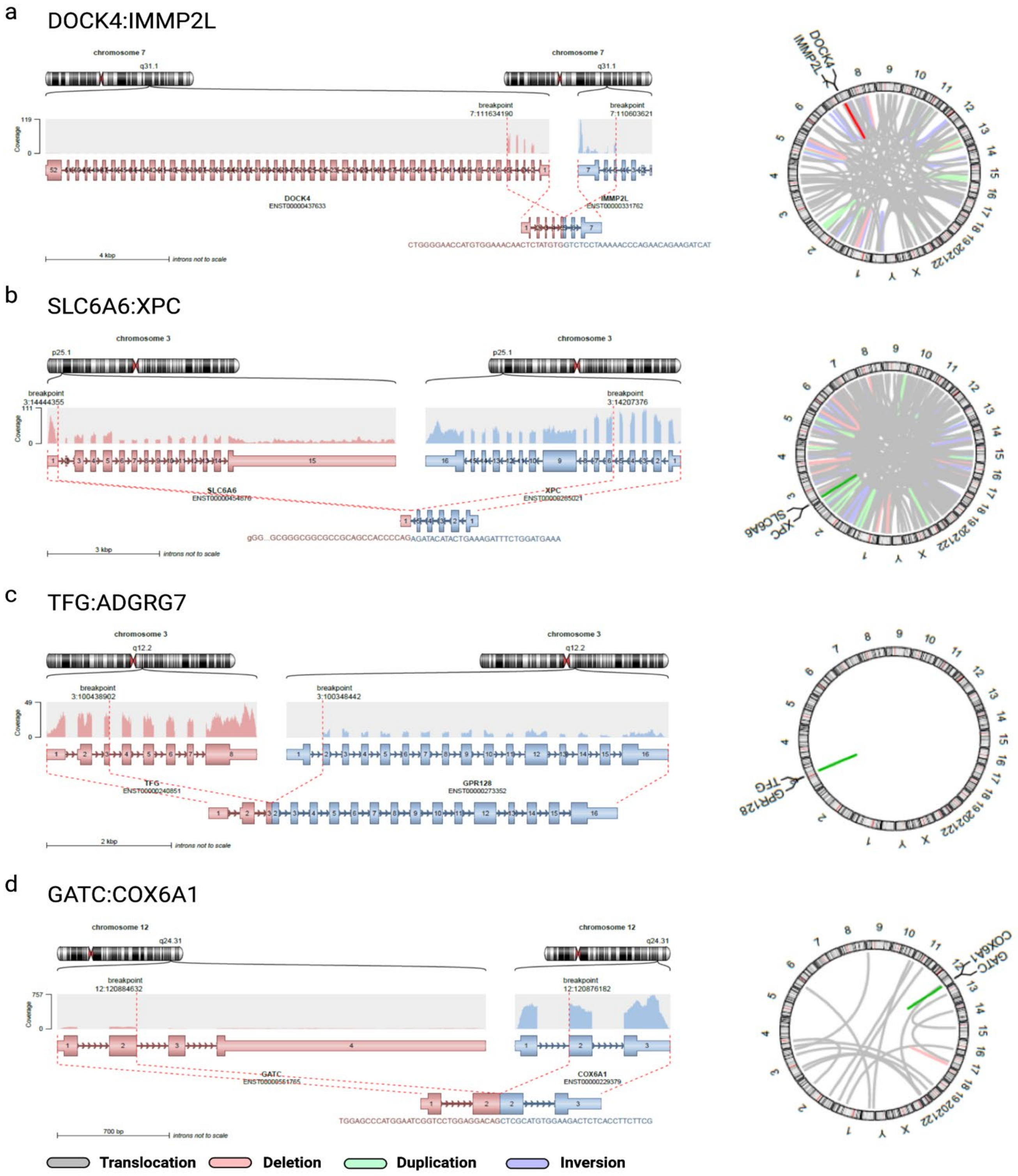


**Figure S5. Prediction of gene fusions in plasma derived EV.** Plasma, paired tumor and healthy controls were analyzed for gene fusions using Arriba, Pizzly and FusionCatcher.Structural representation of gene fusion and circle plot depicting main chromosomal aberrations. **a,** DOCK4:IMMP2L fusion was detected in patient ID 21 and supported by arriba: pair=4, reads=4, pizzly:pair=15, reads=11, and fusioncatcher: pair=7, reads=6. **b,** SLC6A6:XPC and antisense SLC6A6:XPC-AS1 fusions were detected in patient ID 24 and supported by arriba: pair=13,reads=24 and fusioncatcher: pair=16,reads=7 respectively. **c,** TFG:ADGRG7 fusion was detected in patient ID 27 and was supported by fusioncatcher: pair=2, reads=4 and pizzly: pair=1, reads=3. **d,** GATC:COX6A1 was detected in patients IDs 11 and 40 and was supported by arriba: pair =3, reads =2 and arriba: pair=7, reads =26, respectively.
